# Evaluating a Human/Machine Interface with Redundant Motor Modalities for Trajectory-Tracking

**DOI:** 10.1101/2022.06.29.498180

**Authors:** Amber H.Y. Chou, Momona Yamagami, Samuel A. Burden

## Abstract

In human/machine interfaces (HMI), humans can interact with dynamic machines through a variety of sensory and motor modalities. Redundant motor modalities are known to have advantages in both human sensorimotor control and human-computer interaction: motor redundancy in sensorimotor control provides abundant solutions to achieve tasks; and incorporating diverse features from different modalities has improved the performance of movement-, gesture-, and brain-controlled computer interfaces. Our objective is to investigate whether redundant motor modalities enhance performance for a continuous trajectory-tracking task. We designed a multimodal human/machine interface with combined *manual* (joystick) and *muscle* (surface electromyography, sEMG) inputs and evaluated its closed-loop performance for tracking trajectories through second-order machine dynamics. In a human subjects experiment with 15 participants, we found that the multimodal interface outperformed the manual-only interface while performing comparably to the muscle-only interface; and that the multimodal interface enabled users to coordinate individual modalities to attenuate noise. Multimodal human/machine interfaces could be beneficial in systems that require stability and robustness against perturbations such as motor rehabilitation and robotic manipulation.

## 1. INTRODUCTION

In human/machine interfaces (HMI), humans interact and communicate with dynamic machines through one or more *sensory* and *motor* modality. *Sensory* modalities enable humans to perceive stimuli from machines (e.g., a computer display that provides visual stimuli to human eyes), while *motor* modalities enable humans to provide motor inputs to machines (e.g., a joystick that measures hand movement on one or more axes). We refer to traditional hand-held devices such as joysticks, mice, and steering wheels as *manual* modalities.

Recently, researchers have begun implementing biological signals like muscle activity as an alternative motor input. For instance, electromyographic (EMG) signals are widely used in developing body-machine interfaces (Casadio et al., 2012) and to control assistive robotic devices (Kiguchi and Hayashi, 2012; Fall et al., 2017). We refer to the recordings of muscle inputs in HMI as *muscle* modalities in this paper. Both muscle and manual modalities have unique advantages as human interfaces. For instance, muscle interfaces provide high-density measurements (Drost et al., 2006) and have large control bandwidth (Lobo-Prat et al., 2014; Yamagami et al., 2020). However, there are still many challenges in current upper-limb muscle inter-faces such as high signal variability between and within individuals, and high sensitivity to perturbations (Farina et al., 2014; Artemiadis, 2012). On the other hand, manual interfaces have lower control bandwidth but provide less variable signals and are currently much more familiar to users. Fusing the inputs from both motor modalities has the potential to combine benefits and compensate for weaknesses from both sides (Pantic and Rothkrantz, 2003; Rizzoglio et al., 2020).

To develop human/machine interfaces with redundant motor modalities, it is instructive to first understand how humans and other animals integrate motor pathways in their sensorimotor systems. Previously, researchers have assessed the integration of parallel *sensory* pathways (Roth et al., 2016; Peterka, 2018) and *motor* pathways (Gelfand and Latash, 1998; Scholz and Schöner, 1999; Latash et al., 2002) to explain how redundant inputs collectively govern a single behavior. Researchers have also investigated motor redundancy using body-machine interfaces (Ranganathan et al., 2014; De Santis and Mussa-Ivaldi, 2020), in which body inputs were collected from multiple channels with the same modality to control lower-dimensional systems. Studies have found that sensorimotor redundancy could provide abundant solutions to achieve tasks (Todorov, 2004), provide stability (Latash et al., 2002), assure robustness against uncertainties (Gelfand and Latash, 1998; Roth et al., 2016), and enhance movement efficiency (De Santis and Mussa-Ivaldi, 2020).

The advantages of multimodal interfaces have been suggested in previous studies in the fields of human- and brain-computer interfaces. Multimodal interfaces can incorporate diverse features from different modalities, and thus enable human-computer interfaces to better analyze complex human behaviors (Pantic and Rothkrantz, 2003; Jaimes and Sebe, 2007), recognize hand gestures (Wu et al., 2016), and classify intent of movement (Zhang et al., 2019). Similarly, fusing multiple data sources with different frequency bands could enhance the accuracy of brain decoders (Fazli et al., 2015) and improve usability of brain-computer interfaces (Müller-Putz et al., 2011). Most relevant for the present study is the muscle-and-motion interface studied in Rizzoglio et al. (2020), where it was found that the multimodal (*hybrid*) interface performed similarly to motion-only (implemented with inertial measurement units, IMU) and outperformed muscle-only (implemented with surface electromyography, sEMG) in a reaching task.

The objective of this study is to evaluate a human/machine interface that combines redundant muscle and manual motor modalities to perform a second-order reference-tracking and disturbance-rejection task. Our experimental results with 15 human subjects demonstrate that the multimodal interface outperforms the traditional manual interface and performs comparably to the muscle interface. We also found that users can suppress the effect of sensorimotor noise in task-relevant dimensions by coordinating noise between muscle and manual modalities. Our results demonstrate potential advantages for human/machine interfaces that incorporate motor redundancy.

## 2. MATERIALS AND METHODS

We developed a multimodal interface with manual and muscle modalities. We then conducted human-subject experiments and data analyses to empirically evaluate performance of the multimodal interface.

In the following sections, we denote a signal and a transfer function as *q* and *T* in time-domain; and 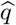 and 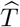 in frequency-domain. The frequency-domain representations of signals are calculated using the fast Fourier Transform (FFT).

### 2.1 Human/Machine System Development

#### Task Development

We adopted a 1-degree-of-freedom (DOF) continuous human/machine task previously conducted by Yamagami et al. (2021). Human users *H* are tasked with controlling a second-order linear-time invariant (LTI) machine dynamics *M* :

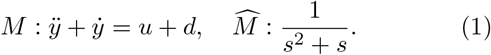

The objective of human users is to control their cursor output *y* to track a reference trajectory *r* on a computer screen and reject the invisible input disturbance *d*. Users control the system using both muscle and manual modalities through a custom interface that records motor inputs. We define the muscle and manual input signals as *u_μ_* and *u_ν_*, respectively. The two inputs were linearly combined with a weight constant, *α*:

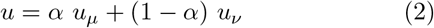

To investigate the optimal weighting of the two modalities, we selected *α* to be evenly spaced numbers in [0, 1], where *α* ∈ {0, 0.25, 0.5, 0.75, 1}. We refer to the weighted sum *u* as the overall user input, and the transformation between tracking error *r* − *y* and *u* as the feedback controller *B*. A block diagram of the human/machine system is shown in Figure 1.

**Fig. 1.**
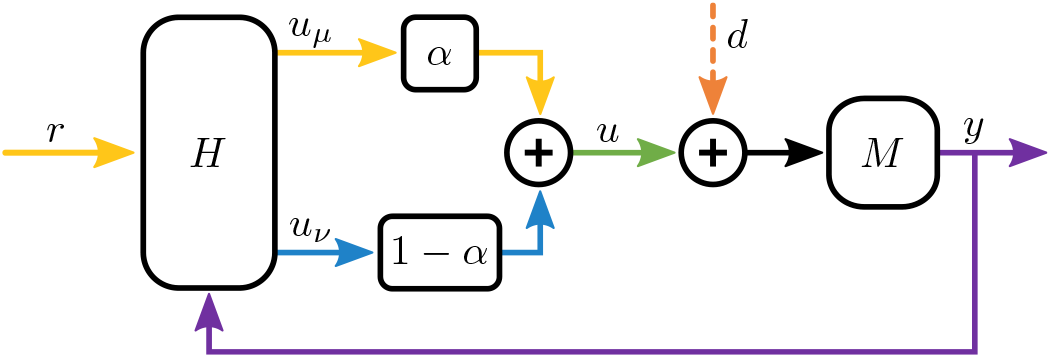
Block diagram of human/machine interface. The human user *H* transforms reference *r* and output *y* to produce *muscle u_μ_* and *manual u_ν_* signals that are scaled and summed to yield user input *u*. User input *u* is added to disturbance *d* and transformed through a second-order machine *M* to produce output *y*.

#### Generating Stimuli

We constructed pseudorandom reference *r* and disturbance *d* as sums of sinusoidal signals interleaved at the stimulated frequencies (Yamagami et al., 2021; Yu et al., 2014). Stimulated frequencies Ω were designed to be eight prime multiples of the base frequency (0.05 Hz) below 1 Hz, where Ω = {0.1, 0.15, 0.25, 0.35, 0.55, 0.65, 0.85, 0.95 Hz}. The magnitude of each stimulus was scaled by the inverse of its frequency, and the phase was randomized.

#### Multimodal Interface Development

We developed a combined muscle and manual multimodal interface for this study (Fig. 2). The manual signals were measured using a 1-DOF 10 *k*Ω slide potentiometer. The slider’s handle has a travel distance of 11.4 cm. The muscle signals were obtained via surface EMG electrodes (Muscle SpikerShields, Backyard Brains, Inc.). The EMG electrodes were placed on participants’ biceps and triceps according to the guidelines of Surface Electromyography for the Non-Invasive Assessment of Muscle (SENIAM) (Hermens et al., 2000).

**Fig. 2.**
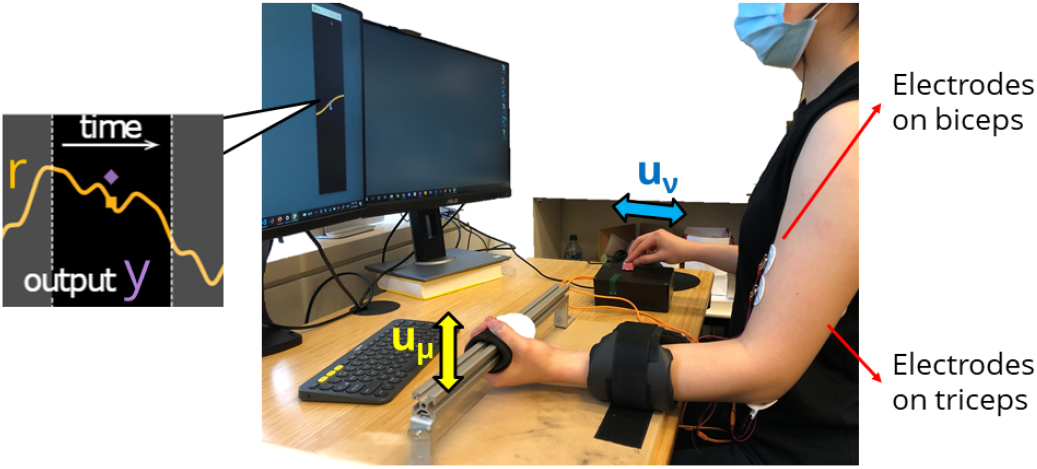
Multimodal human/machine interface: reference trajectory *r* and cursor output *y* are displayed on a computer screen; participants used their non-dominant arm to control the muscle modality (left arm in this image) and the dominant hand to control the manual modality (right hand in this image). Electrodes were placed on participants’ biceps and triceps.

The potentiometer values and the EMG data were recorded with an Arduino Uno (Arduino.cc) at 1000 Hz. We converted the raw EMG data to the Average Rectified Value (ARV) by taking the absolute value of raw EMG signals and applying a moving average filter with a 200 ms window. We then normalized the ARV with the maximum voluntary contraction (MVC) of each participant. To prevent muscle fatigue, only a maximum of 33% of the MVC was required to track the largest stimuli of the task. We then collected the time-domain reference *r*, disturbance *d*, cursor output *y* and user inputs (*u*, *u_μ_* and *u_ν_*) of every trial at 60 Hz sampling rate for later analysis.

### 2.2 Experimental Design

#### Participants

We recruited 15 participants (6 female, 9 male; 14 right-handed, 1 left-handed; aged between 22 to 31 years; height: 177±15.25 cm; weight: 71.5±19 kg). All participants use computers and smartphones daily. Six participants have seen or used EMG devices, and two participants work with EMG regularly.

#### Experimental Setup and Protocol

Participants were asked to perform the task with both of their upper limbs: manipulating the slider with their dominant hand and activating the biceps or triceps of their non-dominant arm. Each participant performed three sessions in the following order:

(1^*st*^ session) 10 trials for each single-mode condition (manual-only *α* = 0, followed by muscle-only *α* = 1);

(2^*nd*^ session) 14 trials for each multimodal condition (*α* = 0.25, 0.5, 0.75 in random order);

(3^*rd*^ session) 4 trials for each single-mode condition (*α* = 0, 1 in random order) as the control groups.

We started each experiment with the manual interface (*α* = 0) to allow participants to familiarize the task since most computer users are more familiar with a manual device. Each trial was 45 seconds starting with a 5 seconds ramp-up period (ramp-up was not included in the data analysis), thus the total time period *T* = 40 *s*. Participants were asked to take mandatory breaks in-between conditions to avoid muscle and eye fatigue.

### 2.3 Hypotheses and Data Analysis

We hypothesized that multimodal interfaces would perform better than single-mode interfaces. We assessed performance of interfaces using (1) overall tracking performance, (2) reference-tracking performance at stimulus frequencies, (3) disturbance-rejection performance at stimulus frequencies, and (4) user response at non-stimulated frequencies, which we regard as sensorimotor noise.

#### Hypothesis 1

*Tracking performance is higher in multi-modal conditions than single-mode conditions.*

We quantified the overall task performance using the time-domain mean-square error (MSE) between reference (*r*) and cursor output (*y*):

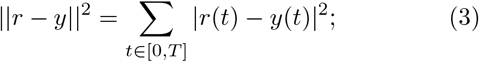

lower error corresponds to better performance.

#### Hypothesis 2

*Reference-tracking performance at stimulus frequencies is higher in multimodal conditions than single-mode conditions.*

Given a LTI machine *M* and stimuli at specific frequencies, we can assume that humans *H* behave approximately like an LTI transformation (Yamagami et al., 2021). Hence the output 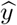 can be written as linear combination in response to reference 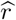 and disturbance in the output space 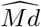:

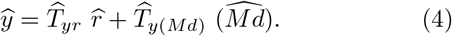

The user response to reference stimuli can be investigated by calculating the system-level transformation 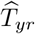. Since reference and disturbance stimuli are interleaved at the stimulated frequencies, when 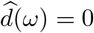,

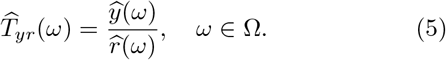

In a perfect reference tracking scenario, 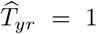 with gain 1 and phase 0. We quantified the reference tracking performance as error between the transformation and 1:

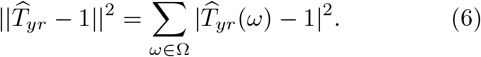

We scaled 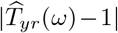 by the magnitudes of output stimuli in order to emphasize the effect of tracking errors in the lower frequencies.

#### Hypothesis 3

*Disturbance-rejection performance at stimulus frequencies is higher in multimodal conditions than single-mode conditions.*

Given (4), when 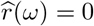, the user response to disturbance stimuli can be investigated by calculating the system-level transformation 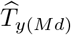:

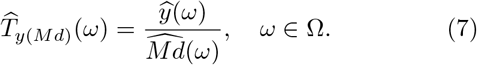

If the user can perfectly reject disturbance in the output space, 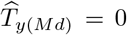 with gain 0. Therefore, we quantified the disturbance rejecting performance as error between the transformation and 0:

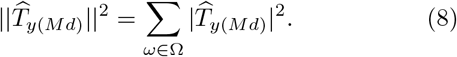

Similarly to (6), the transformation was scaled by the magnitudes of output stimuli.

#### Hypothesis 4

*Sensorimotor noise is lower in multimodal conditions than single-mode conditions.*

In addition to the analyses on performance in tracking/rejecting the stimuli, we analyzed the human sensorimotor noise. To compute the human sensorimotor noise as the imputed disturbance 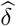, we first estimated the feedback controller 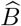 (Yamagami et al., 2021):

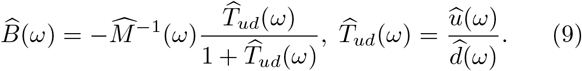

We then defined the open loop transfer function 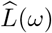:

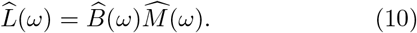

We defined the human sensorimotor noise as the imputed disturbance 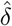, which is the input disturbance if the effect of reference and disturbance stimuli are removed in the feedback system:

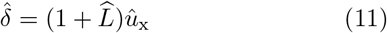

where 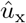 is the user input at the non-stimulated frequencies below 1 Hz. To find 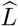 at non-stimulated frequencies, we linearly interpolated the open-loop transfer function values at stimulated frequencies, 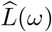, since the transfer function can only be computed at stimulated frequencies.

#### Statistical Tests

To verify the four hypotheses above, we considered the average of the last four trials of each condition in the 2^*nd*^ and the 3^*rd*^ sessions to ensure adequate user learning time. we compared each of the three multimodal conditions (2^*nd*^ session) with each single-mode condition (3^*rd*^ session) by applying Wilcoxon signed-rank test (non-parametric paired t-test) with a confidence level of 5%.

## 3. RESULTS

### 3.1 Multimodal Interface Enhanced Tracking Performance

We found that the equal-weighted interface improved overall performance compared to the manual interface, but not the muscle interface. Time-domain MSE of the equal-weighted condition (*α* = 0.5) was significantly lower than the manual-only condition (*α* = 0) (Wilcoxon signed-rank test: *p* < 0.05) (Fig. 3a). Moreover, time-domain MSE of the muscle-dominated condition (*α* = 0.75) had no difference with the manual-only condition but was significantly higher than the muscle-only condition (*α* = 1) (Fig. 3a). These findings partially supported our **Hypothesis 1**.

**Fig. 3.**
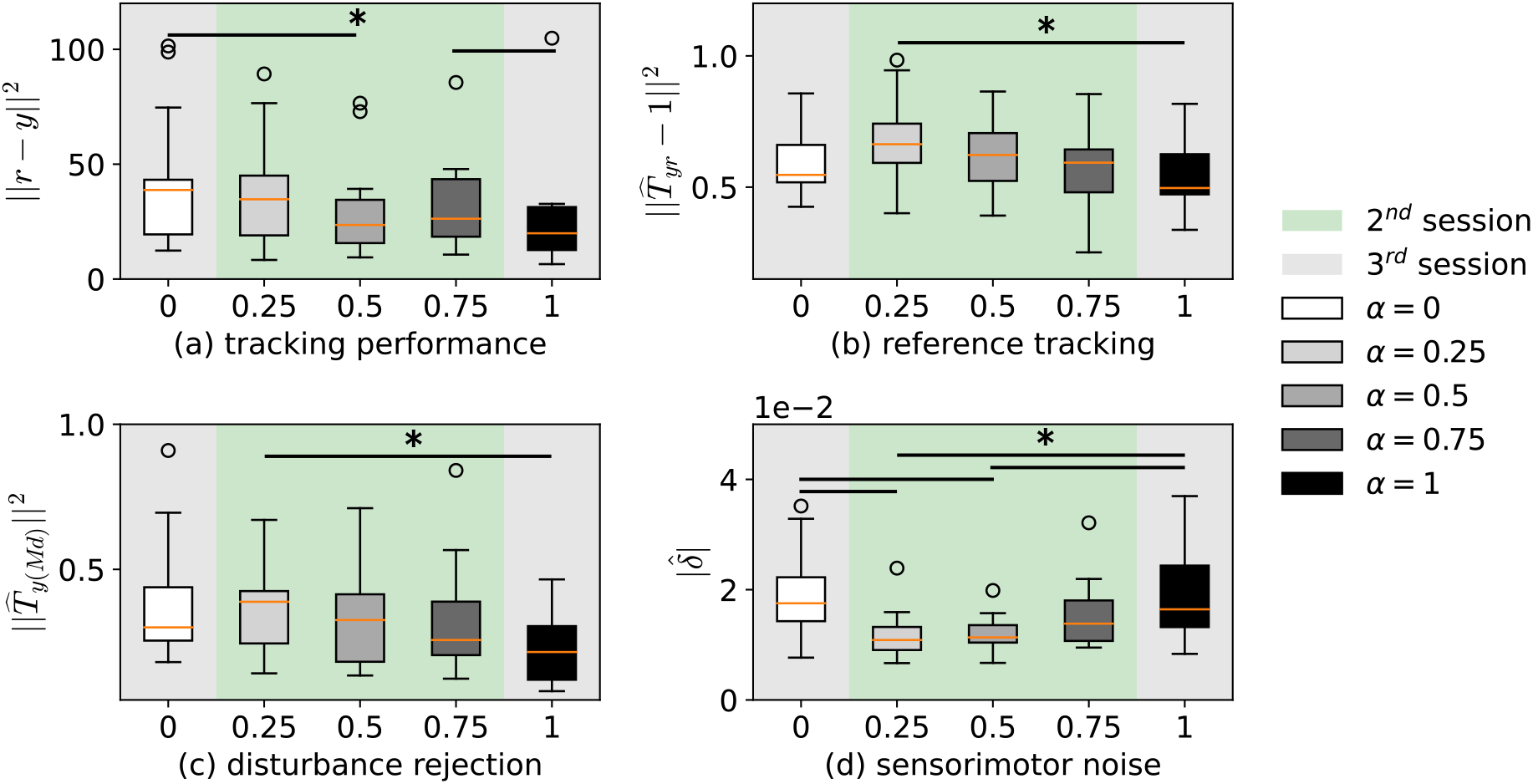
Distributions (median, interquartile) of task performance of the five conditions in the latter two sessions: (a) tracking performance (time-domain MSE); (b) reference-tracking performance; (c) disturbance-rejection performance; and (d) sensorimotor noise. Horizontal lines indicate significant differences between pairs of distributions (Wilcoxon signed-rank test, \**p* < 0.05).

### 3.2 Multimodal Interface Did Not Improve Reference Tracking or Disturbance Rejection

We then tested whether the improvement of the equal-weighted interface was due to better tracking or rejecting of stimuli. However, we did not observe significant difference between the equal-weighted condition and the single-mode conditions in 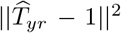 (Fig. 3b) or 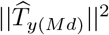 (Fig. 3c). In addition, the manual-dominated condition (*α* = 0.25) performed significantly worse in both reference tracking and disturbance rejection than the muscle-only condition (*α* = 1). These findings led us to reject our **Hypothesis 2** and **3**.

### 3.3 Multimodal Interface Reduced Sensorimotor Noise

We then investigated whether the improvement in time-domain performance was due to better noise suppression. We found that the equal-weighted (*α* = 0.5) and the manual-dominated condition (*α* = 0.25) had significantly lower (*p* < 0.05) imputed disturbances *δ* than both the single-modal conditions (Fig. 3d). We did not observe a significant noise reduction in the muscle-dominated (*α* = 0.75) interface (Fig. 3d). These findings partially supported our **Hypothesis 4**.

### 3.4 Multi-Modal User Inputs Were Anti-Correlated at Non-Stimulated Frequencies

We further investigated the user response at non-stimulated frequencies, *u*_x_, to understand how humans suppressed sensorimotor noise in multimodal conditions. We denoted the inputs from each modality at the non-stimulated frequencies as 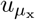 and 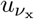. We found that humans combined both modalities antagonistically at the non-stimulated frequencies. For all multimodal conditions, the magnitudes of overall 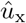 were the weighted combination of 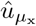 and 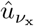 above crossover frequency^1^ (Fig. 4). However, 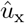 had lower magnitude than 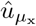 and 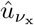 below the crossover frequency. The differences in phases, 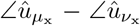, were approximately *π* below crossover (Fig. 4).

**Fig. 4.**
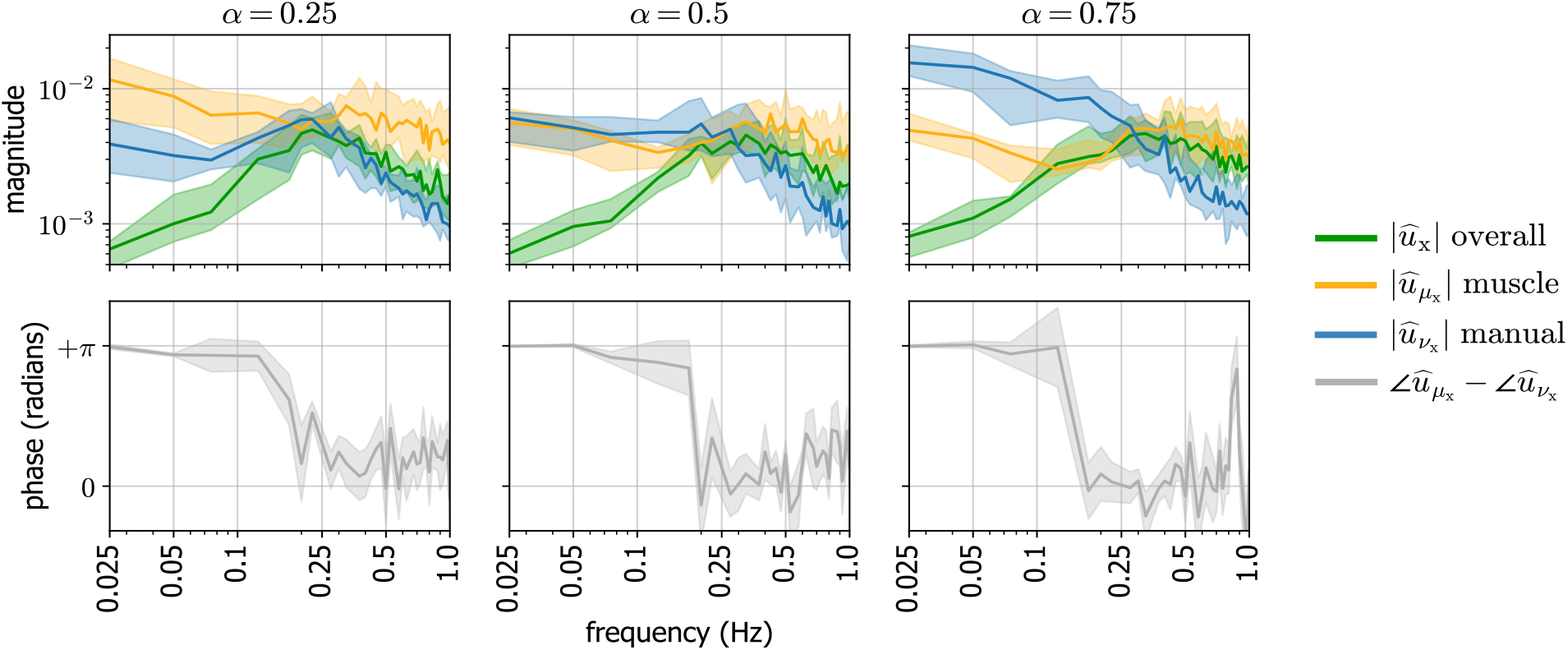
Distributions (median, interquartile) of user responses at non-stimulated frequencies in multimodal conditions. (*top*) Magnitudes of overall input (*u*_x_) and inputs from individual two modalities 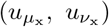. (*bottom*) Phase difference between 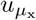 and 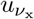.

## 4. DISCUSSION

We found that an equal-weighted multimodal interface outperformed a manual-only interface due to reduced sensorimotor noise, and had comparable performance to a muscle-only interface. This observation is similar to the finding of Rizzoglio et al. (2020), where a multimodal interface outperformed one single-mode interface and had comparable performance to another. However, our results differ in that we found performance improvements for a multimodal interface relative to a *manual* but not *muscle* modality, whereas the previous study of Rizzoglio et al. (2020) found benefits for multimodality relative to a *muscle* but not *motion* (IMU) modality – which is related to, but distinct from, our *manual* (slider) modality.

The improvement in the equal-weighted multimodal interface did not extend to our unequal-weighted interfaces. The muscle-dominated condition performed significantly worse in time-domain tracking than the muscle-only condition, while the manual-dominated condition had significantly higher error in tracking/rejecting stimuli compared to the muscle-only condition. We additionally found that the equal-weighted and the manual-dominated interfaces enhanced sensorimotor noise suppression compared to both single-mode interfaces. This observation in noise suppression aligns with prior findings that sensorimotor redundancy assures steady motions and robustness against perturbations (Gelfand and Latash, 1998; Latash et al., 2002; Roth et al., 2016).

Our investigation in user inputs at non-stimulated frequencies show that humans had control over both *magnitudes* and *phases* of their inputs for frequencies below crossover. Humans increased the *magnitude* of their inputs if there was a “shortage” in a modality. For instance, in the manual-dominated condition, users increased 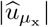 below crossover to compensate the lower weighting in muscle input (Fig. 4). In addition, users coordinate the non-stimulated signals of the two modalities in opposite *phases* below crossover. If the noise signals on different modalities are independent, the signals would not combine constructively. This suggests that the integration of motor pathways attenuates sensorimotor noise. We think our finding in opposite phases of the redundant modalities is similar to the Uncontrolled Manifold concept (Scholz and Schöner, 1999; Todorov, 2004). Future work may investigate how noise varies over longer periods of time (Huber et al., 2016) and, more broadly, the role of noise in skill acquisition (Sternad, 2018).

## 5. CONCLUSION

In this study, we suggested an experimental method to evaluate multimodal human/machine interfaces for a second-order continuous trajectory-tracking and disturbance rejection task. Experimental results demonstrated that the equal-weighted interface enhanced the time-domain performance compared to the manual interface; in addition, the equal-weighted interface had comparable performance to the muscle interface but had a significantly lower sensorimotor noise. We also investigated the user responses at non-stimuli and found that humans had control over both the magnitudes and phases of each modality at low frequencies. Moreover, the antagonistic phases of the two modalities led to lower sensorimotor noise of the multimodal interfaces. These observations provide evidence that multimodal interfaces can better suppress sensorimotor noise through redundant motor modalities. Our results suggest that future human/machine systems that require minimum sensorimotor noise may benefit from redundant motor modalities, especially combined manual and muscle modalities.

## ACKNOWLEDGEMENTS

We thank the participants in this study, and our colleague Joshua Vasquez for manufacturing assistance.

## Footnotes

1 frequency at which the open-loop transfer function magnitude is below 1, 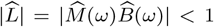 (McRuer and Jex, 1967). For our dataset, the crossover frequency was computed to be around 0.25 Hz.

